# In vivo structure and dynamics of the RNA genome of SARS-Cov-2

**DOI:** 10.1101/2021.01.15.426526

**Authors:** Yan Zhang, Kun Huang, Dejian Xie, Jian You Lau, Wenlong Shen, Ping Li, Dong Wang, Zhong Zou, Shu Shi, Hongguang Ren, Meilin Jin, Grzegorz Kudla, Zhihu Zhao

## Abstract

The SARS-CoV-2 coronavirus, which causes the COVID-19 pandemic, is one of the largest positive strand RNA viruses. Here we developed a simplified SPLASH assay and comprehensively mapped the in vivo RNA-RNA interactome of SARS-CoV-2 RNA during the viral life cycle. We observed canonical and alternative structures including `3’`-UTR and `5’`-UTR, frameshifting element (FSE) pseudoknot and genome cyclization in cells and in virions. We provide direct evidence of interactions between Transcription Regulating Sequences (TRS-L and TRS-Bs), which facilitate discontinuous transcription. In addition, we reveal alternative short and long distance arches around FSE, forming a “high-order pseudoknot” embedding FSE, which might help ribosome stalling at frameshift sites. More importantly, we found that within virions, while SARS-CoV-2 genome RNA undergoes intensive compaction, genome cyclization is weakened and genome domains remain stable. Our data provides a structural basis for the regulation of replication, discontinuous transcription and translational frameshifting, describes dynamics of RNA structures during life cycle of SARS-CoV-2, and will help to develop antiviral strategies.

## Introduction

The COVID-19 pandemic, caused by the SARS-CoV-2 coronavirus, has caused more than 1.5 million deaths worldwide up till the time of submission. Although many efforts have been devoted to control the disease, vaccines are being approved for emergency use, but the pandemic is far from being under control. Therefore, there is an urgent need to understand basic molecular biology of SARS-CoV-2 coronavirus.

SARS-CoV-2 belongs to the broad family of coronaviruses. It is a positive-sense single-stranded RNA virus, with a single linear RNA segment of approximately 30,000 bases [1].

Coronavirus RNA-dependent RNA synthesis includes two different processes: continuous genome replication that yields multiple copies of genomic RNA (gRNA), and discontinuous transcription of a collection of subgenomic mRNAs (sgRNAs, or sgmRNAs) that encode the viral structural and accessory proteins [2, 3]. The transcription process is controlled by transcription-regulating sequences (TRSs) located at the 3′ end of the leader sequence (TRS-L) and preceding each viral gene (TRS-B), and requires base-pairing between the core sequence of TRS-L (CS-L) and the nascent minus strand complementary to each CS-B (cCS-B), allowing for leader-body joining [2, 4, 5]. A three-step working model of coronavirus transcription was suggested [2, 6], which implies that long-distance RNA-RNA interactions are required prior to template switch. Long-distance interactions between B motif (B-M) and its complementary motif (cB-M), and between proximal element (pE) and distal element (dE), are important for forming high-order structures promoting discontinuous RNA synthesis during N sgmRNA transcription in the TGEV coronavirus [5]. However, the motifs involved in these interactions are not conserved in beta-coronaviruses, and it is not known if similar interactions contribute to transcription of other sgmRNAs. Therefore, although it is widely assumed that TRS-L interacts with cCS-B, there has been no experimental evidence for direct interactions between TRS-L and TRS-Bs.

Functional studies have revealed the importance of RNA secondary structures for viral replication, transcription and translation [7-9]. One unique feature of the coronavirus is frameshifting in ORF1ab, giving rise to RNA-dependent RNA polymerase (RdRP) and other proteins in ORF1b. The structure of the SARS-CoV FSE (whose sequence differs from the SARS-CoV-2 FSE by just one nucleotide) was solved by NMR to be a three-stem pseudoknot [10], and was supposed to play key roles in translational control of ORF1b [11]. Genome-wide strategies to chemically probe RNA structure in cells made in vivo SARS-CoV-2 RNA structure analysis available [12-15]. Recently, alternative conformations of the frameshift element (FSE) were derived from in-cell SHAPE/DMS-MaP seq data [14, 15]. Although these methods provided insight into cis-acting RNA structures regulating important biological processes of virus life cycle, they could not elucidate long distance interactions. Additionally, short- and long-distance interactions within the SARS-CoV-2 RNA were described using the COMRADES and vRIC-seq methods. Importantly, Ziv et al. discovered networks of both genomic RNA (gRNA) and subgenomic RNA (sgRNA) interactions by applying specific probes to pull down each RNA species, and Cao et al reconstructed structures in virions[16]. However, all of these experiments were performed in a specific stage of the virus life cycle. So, there is need to directly compare structures from different stages to investigate their dynamics and functional relevance during the whole life cycle.

In this study, to comprehensively map RNA-RNA interactions of SARS-CoV-2 RNA both in cells and in virions, we simplified sequencing of psoralen crosslinked, ligated, and selected hybrids (SPLASH) [17] based on proximity ligation, with three major differences: 1) RNase III was used to fragment RNA as in PARIS [18], this treatment makes fragmented RNA ends compatible for T4 RNA ligase, 2) T4 PNK treatment is omitted, 3) the purified ligation products were directly subjected to a commercial pico-input strand-specific RNA-seq library construction kit. The major purpose of these modifications is to make the protocol more suitable for low amount of virion RNAs. To investigate the dynamics in RNA structure during life cycle of SARS-CoV-2 virus, we performed simplified SPLASH on early, late infected cells and supernatant virions.

Here we provide direct experimental evidence of comprehensive TRS-L interactions with TRS-B regions of sgRNAs, and identify and validate novel sgRNAs by analyzing additional TRS-L interaction peaks. We found multiple alternative interactions mediated by FSE, providing structural basis for ribosome stalling. In addition, we showed that both proximal and distal genome RNA-RNA interactions are strengthened, while sgRNAs mediated interactions are significantly reduced in virions, suggesting thorough compaction of genome RNA in virions. Interestingly, although TRS-L mediated interactions including genome cyclization are weakened, interactions between TRS-L and ORF S are strengthened in later phase of infection cells and mature virions, which may contribute to rapid transcription of sgRNAs. Our data provides a comprehensive overview of relationships between SARS-CoV-2 RNA structure and key processes of virus life cycle, such as replication, discontinuous transcription and translation.

## Results

### Overview of short- and long-range RNA-RNA interactions of SARS-Cov-2

We developed a simplified SPLASH protocol to capture RNA-RNA interactions in the SARS-CoV-2 virus. Briefly, the biological samples are first stabilized by Psoralen-PEG3-Biotin cross-linking, followed by RNase III treatment, proximity ligation, library preparation, and high throughput sequencing. Samples were collected from different phases of the SARS-CoV-2 virus life cycle to infer the dynamic structure of viral RNA at different stages. In the early stage of infection, we collected virus-infected Vero cells (C), in which cytopathic effect (CPE) was not observed. At a later stage, when 70% of cells underwent CPE, we collected the cell culture supernatant and harvested mature virus particles (V) and, at the same time used freeze-thaw methods to lyse the cells (L) to collect the cell and virus RNA (Figure 1A). The major steps and corresponding RNA are shown in Figure 1A. The ligated RNA fragments could form both `5’`-`3’` and `3’`-`5’` chimeras [19] (Figure 1C and Figure 1D). Pearson correlation analysis of chimera counts between biological replicates indicated high reproducibly of experiments (Figure 1B and Supplementary Figure S1A and S1B). Chimeric signals were significantly higher in ligated than in non-ligated samples (Figure 1C and supplementary table 1) As documented by others [20, 21], we also observed weak chimeric signals in non-ligated samples. Counts of chimeras in ligated and non-ligated samples were correlated, particularly in the C sample (Figure S1C∼E), consistent with the idea that chimeras in non-ligated samples may due to endogenous ligation activity derived from host cells.

**Figure 1.**
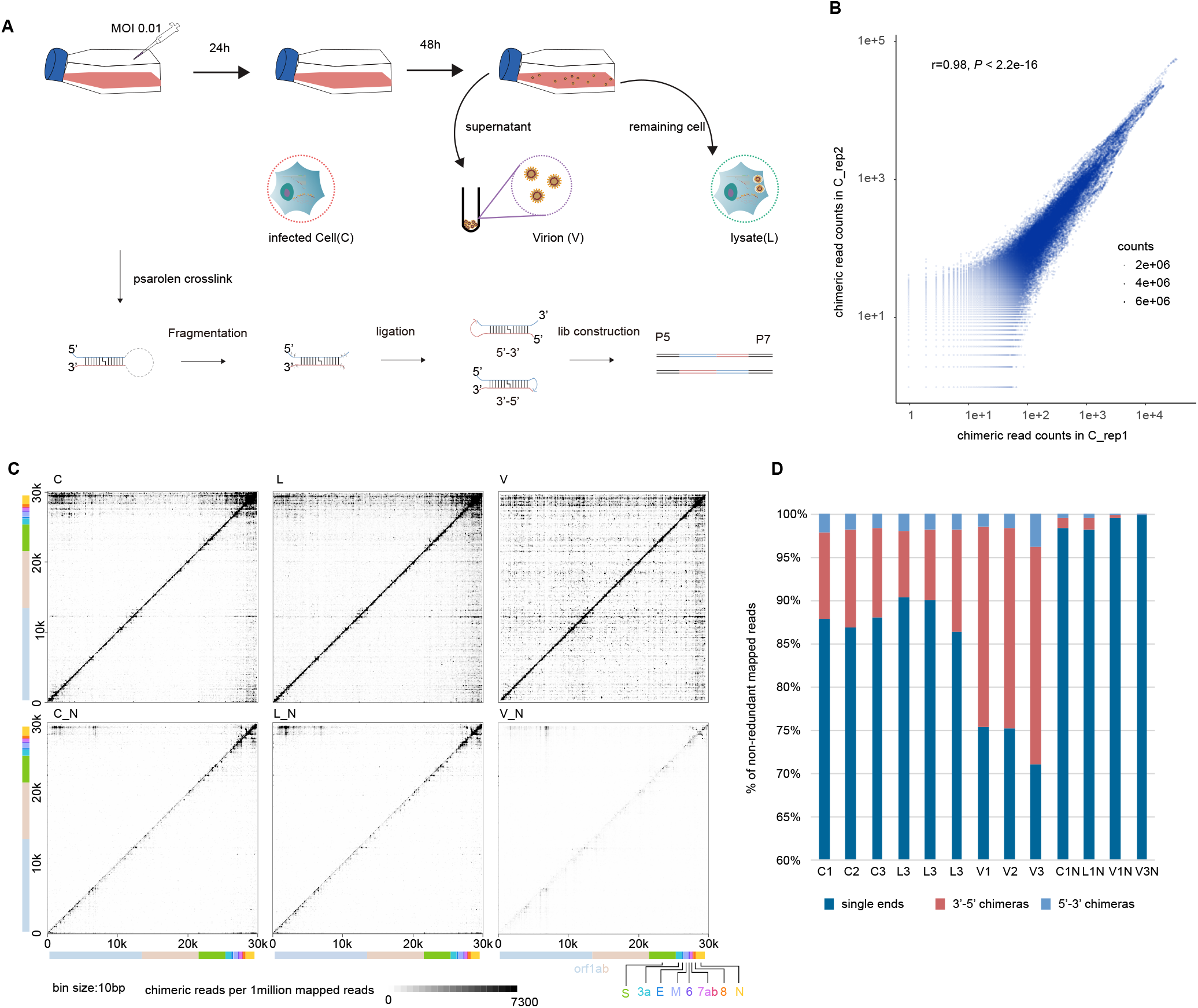
Overview of the experiment. (A) Schematic diagram for sample collection and major experimental steps. (B) Dotplot shows chimeric read counts from two replicates, indicating good reproducibility of proximity protocol. (C) Heat map of RNA-RNA interactions along the SARS-CoV-2 gRNA. Each dot represents an interaction signal between the genomic coordinates on the x and y axes. X axis shows the coordinates of the 5’ arm of the chimera, and the Y axis shows the 3’ arm of the chimera. So, 5’-3’ chimeras are above the diagonal, and 3’-5’ chimeras are below the diagonal. (D) Statistics of mapped single end RNA, `3’`-`5’` chimeras and `5’`-`3’` chimeras in each sample.

### Structures of UTRs and genome cyclization

The `5’`-UTRs of coronaviruses contain five evolutionarily conserved stem-loop structures (denoted SL1–SL5) that are essential for genome replication and discontinuous transcription [8, 22, 23]. In our data, all five stems are supported by chimeras (Figure S2A and S2B). Notably, SL1∼SL3 are stronger in cells than in virions, possibly indicating that these stem loops form more alternative structures in cells. (i.e TRS-L: TRS-B, or genome cyclization which will be discussed below). SL4 is weaker(Figure S2A), supporting the notion that SL4 might function as a structural spacer [24].

The `3’`-UTR contains a bulged stem-loop (BSL), hyper-variable region (HVR) comprising a conserved octonucleotide sequence and the stem-loop II-like motif (S2M), which are essential for sgRNA synthesis in MHV [7, 25, 26]. All of these structures could be detected in our data. Notably, the pseudoknot in the `3’`-UTR had fewer chimeras, whereas S2M had strong signal. Therefore, we identified an alternative `3’`-UTR structure in the S2M region (Figure S2C), similar to a recently reported structure [13].

In the contact matrices, we also found interactions formed by `5’`-end and `3’`-end of the SARS-CoV-2 genome, indicating genome cyclization (Figure 1C and Figure S2E and S2F), which was previously described [27]. Interestingly, we found additional base pairing at genome cyclization sites (Figure S2F), which suggests that SL4 in `5’`-UTR is also involved in cyclization processes. Notably, genome cyclization is reduced in virions (Figure S2E). Genome cyclization was also described in other viruses, including flaviviruses [28], and involved in replicase recruitment at least in Dengue virus and Zika virus [9]. Considering dynamics of genome cyclization upon packaging and releasing, we speculate that genome cyclization is involved in replication and/or packaging. Perturbation of genome cyclization might offer an interesting avenue to target SARS-CoV-2 replication.

### Long distance interactions between TRS-L and TRS-B regions

Although RNA proximity ligation generates chimeric reads indicative of RNA-RNA interactions, cells also contain many spliced transcripts, or in the case of coronavirus-infected cells, sgRNAs, that resemble chimeras produced by proximity ligation because they originate from disjoint regions of the genome. To correctly identify RNA-RNA interactions, it is therefore essential to filter chimeras that result from splicing or discontinuous transcription. In previous studies, filtering was performed by mapping reads to a database of known transcripts, and removing reads mapped to known splice junctions [29]Ramani, 2015, 26237516;Lu, 2016, 27180905}. However, SARS-CoV-2 produces an extreme diversity of sgRNAs [3], many of which are not yet annotated, rendering this approach impractical. Instead, we empirically assessed the characteristics of chimeric reads found in a published RNA-Seq dataset from SARS-CoV-2-infected cells [3], which we assumed to represent sgRNAs rather than RNA-RNA interactions.

We identified three characteristics that differentiated most sgRNA chimeras found in RNA-Seq from bona-fide proximity ligation chimeras: (1) sgRNAs were ligated almost exclusively in the 5`’`-3`’` orientation (i.e. the 5`’`-proximal fragment of the genome corresponded to the 5`’`-proximal fragment of the chimeric read), whereas RNA proximity ligation chimeras can be ligated in the 5`’`-3`’` orientation (also known as inline, forward, or regularly gapped), and 3`’`-5`’` orientation (also known as inverted, reverse, or chiastic) (reviewed in [19]); (2) the junctions between arms of chimeras were precisely localised in sgRNAs, whereas ligation sites in proximity ligation were variable, due to the random nuclease digestion step used in proximity ligation [30]; (3) sgRNA chimeras typically included regions of homology between TRS-L and TRS-B sides of the chimera [31], whereas proximity ligation chimeras typically include no such regions [30]. Adjustment of the maximum gap/overlap setting in our analysis pipeline, hyb, allows detection (gmax=20, “relaxed pipeline”) or removal (gmax=4, “stringent pipeline”) of most sgRNA chimeras in RNA-Seq data, while proximity ligation chimeras are detected with both settings (Supplementary table1). Notably, chimeras detected with the relaxed pipeline are almost all in 5`’`-3`’`orientation (Figure S3B), and junction sites are highly localised (Figure S3C and S3D). We therefore used the stringent pipeline to analyse proximity ligation data while filtering away contaminating sgRNAs.

In order to identify RNA-RNA interactions mediated by TRS-L, we applied a viewpoint analysis [28] to 5`’`-3`’` and 3`’`-5`’` chimeras. We found multiple TRS-L interaction peaks along the SARS-CoV-2 genome, and these peaks were adjacent to the 5`’`-end of canonical sgRNA regions (Figure 2A, 2B and 2D) and particularly obvious for 3`’`-5`’` chimeras. By contrast, there were few 3`’`-5`’` chimeras in non-ligated samples or RNA-seq data (Figure S3A). To rule out the possibility that these 3`’`-5`’` chimeras come from sgRNAs, we analyzed the distribution of junction sites of TRS-L, and found that these sites were highly variable (Figure 2C). Furthermore, the varied mapped positions of both arms of chimeric reads spanning junction sites, as shown in Figure 2E and S3D for examples, further supported the origin of chimeras from genome folding rather than sgRNA transcripts. Therefore, simplified SPLASH data contains both TRS-L mediated RNA-RNA interactions and TRS-L dependent sgRNAs, which can be discriminated by our methods. RNA base pairing mediated by long range interaction indicated that TRS-L may stably associate with TRS-B regions (Figure 2F and Figure S4). Interestingly, we noticed that TRS-L usually does not interact with the exact TRS-B sequence, but with a flanking sequence within 50nt away. This might provide flexibility for the next step of paring to Ccs-B and template switching.

**Figure 2.**
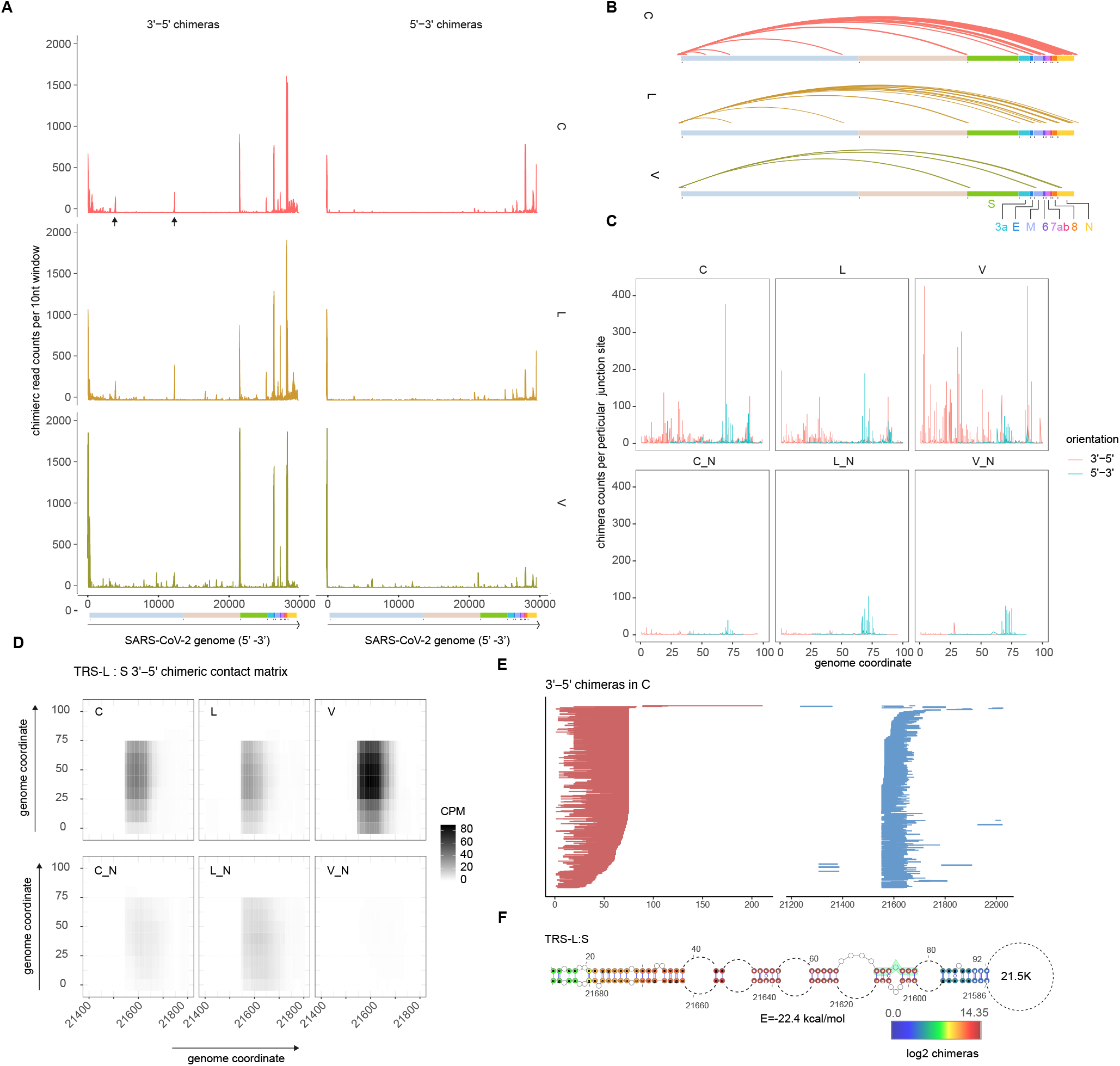
TRS-L interact with canonical TRS-B sites. (A) Viewpoint histograms showing binding positions of the TRS-L region (first 100nt) along the SARS-CoV-2 genome in indicated samples. `3’`-`5’`chimeras and `5’`-`3’` chimeras were separately plotted. Black arrowheads indicate additional peaks in orf1a. (B) Enriched TRS-L interaction peaks deduced from Z-score method. Chimeric read counts from bin-bin contacts were normalized by Z-score, then interactions with Z-score > 2.13 (95% confidence of being above average) and mediated by TRS-L were plotted. (C) Junction site distribution on TRS-L region (the first 100nt), chimeras that break at exactly particular base were counted, showing that the ligation happed in varied sites. (D) Contact matrix of `3’`-`5’`chimeric reads spanning TRS-L: S junction sites. Color depicts counts of chimeric reads per 1million mapped reads (CPM). (E) Randomly selected `3’`-`5’` chimeras overlapping the TRS-L: S junction sites. The red lines indicate `3’`arms of chimeric reads, while blue lines indicate `5’`arms of chimeric reads. Chimeric reads with varied ends are derived from random fragmentation and ligation, reflecting long-range RNA-RNA interactions. (F) RNA base pairing between TRS-L and upstream of S, paired bases were colored by log2 chimeric read counts supporting each base pair (in C sample).

### Identification and validation of novel TRS-L dependent sgRNAs

Apart from canonical sgRNAs, we also observed additional regions interacting with TRS-L (black arrowhead indicated in Figure 2A), with one of them (3.9K) also identified in Ziv’s recent report [27]. The contact matrix based on 3`’`-5`’` chimeric reads and an analysis of individual chimeras showed specific interactions (Figure S5). These regions form stable base pairing with the TRS-L region (Figure S5C and S5E). To check if the TRS-L mediated interactions give rise to new candidate sgRNAs, we performed RT-PCR in independent non-crosslinked cells. Sanger sequencing results confirmed these sgRNAs indeed exist (Figure S5F and S5G). Interestingly, although these novel sgRNA have no canonical ACGAAC core sequence motif (CS-B), they both partially overlap with the canonical CS motif. This indicates that TRS-L and partial cCS-B base pairing at negative strand are critical for template switching in discontinuous transcription of these sgRNAs.

It should be interesting to further validate expression and function of the novel transcripts in the future. This analysis also emphasizes the value of our experiment in dissecting interaction and discontinuous transcription of coronaviruses.

### Alternative structure around frame shift elements

A characteristic feature of coronaviruses is the programmed −1 ribosomal frameshifting to facilitate translation of ORF1b encoding RdRp and controls the relative expression of their proteins. Next, we sought to analyze both local and long range interactions around FSE.

In our data, the proposed three-stem pseudoknot structure [11] was supported by chimeric reads (Figure 3A and 3B), while at the same time, we also found alternative local structures embedding the FSE in larger stable stem loops (arch1), which are all supported by chimeras (Figure 3B, alternative structures 1 and 2). Surprisingly, we also found several alternative long range interactions mediated by FSE (Figure 3A). Besides FSE-arch identified by Ziv et al (referred to as Ziv’s arch hereafter), the alternative arches are formed by FSE and upstream ∼620 nt (arch 2) and ∼1.1 kb (arch 3) elements respectively (Figure 3A). These elements form stable base pairing with FSE (Figure 3B and 3C). 3D structure modeling of the FSE around region (12K-15K) also revealed the spatial proximity between arch 2 and Ziv’s arch (Figure 3D).

**Figure 3.**
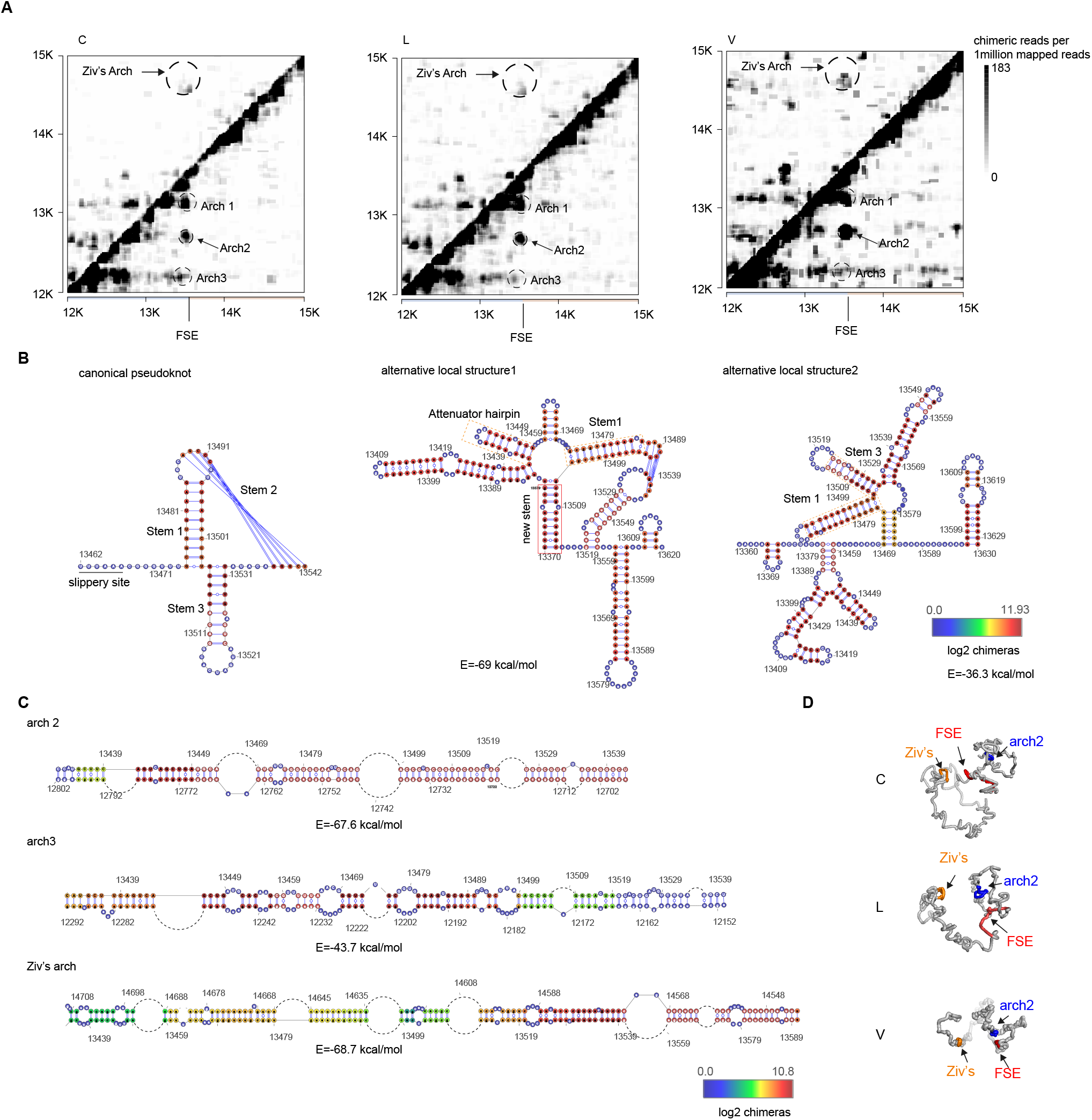
Alternative local and long distance FSE structures. (A) heatmaps show chimeric reads spanning 12K to 15k of SARS-CoV-2 genome in individual samples. Ziv’s arch and alternative arches were plotted as indicated. (B) Basepairing of indicated structures overlaid by log2 chimeric reads in C samples. Left: canonical peudoknot; middle: nt 13370-13620 was folded by minimum energy, and overlaid by log2 chimeric reads in C samples. Right: nt 13360-13630 was folded by COMRADES, and overlaid by log2 chimeric reads in C samples. (C) Basepairing of indicated arches. Colors represent the log2 chimeric read counts of non-redundant chimeric reads supporting each base-pair. (D) 3D modeling of structure around FSE. FSE is in red, while nt 14548-nt14708 (arch4 partner) is orange, and nt 12702-nt12802 is blue.

### Dynamics of RNA structure during viral life cycle of SARS-CoV-2

Next, we analyzed interaction dynamics during phases of viral life cycle. A correlation analysis showed that samples from the same treatment group clustered together (Figure S6A). PCA analysis on chimeric read counts indicates that virion RNA underwent major conformation alteration compared to RNAs in cell (C) and in lysate (L), as shown along primary component (PC) 1 (Figure S6B).

We then used DESeq2 [28, 32] to analyze interactions in each 100 nt × 100 nt window in the viral interaction map. After removal of the low-abundant pairs, pairwise comparisons between C, L and V groups were made. Under a default cutoff (log2FC > ±1, FDR < 0.05), we found similar patterns of differential interactions in the comparisons of virions vs cell (VvsC) and virion vs lysate (VvsL) (Figure S7A), and fold changes of VvsC are correlated with both VvsL and LvsC (Figure S7B). This is concordant with the close relationship between C and L groups in PCA (Figure S6B), suggesting the RNA conformation changes gradually from C to L and then to V.

A heatmap analysis of differential interactions (Figure 4A) suggested a lower density of interactions in the `3’`-third of SARS-CoV-2 genome in virions, compared to cells and lysates. Genome cyclization [28] was also reduced in virions, while proximal interactions and long range interactions other than end-to-end cyclization were strengthened in virions (Figure 4A). An increase in proximal and long-range interactions could also be observed in lysate and still visible when log2FC cutoff was elevated to 5 (Figure S7C). This indicated compaction of genome during packaging into virions.

**Figure 4.**
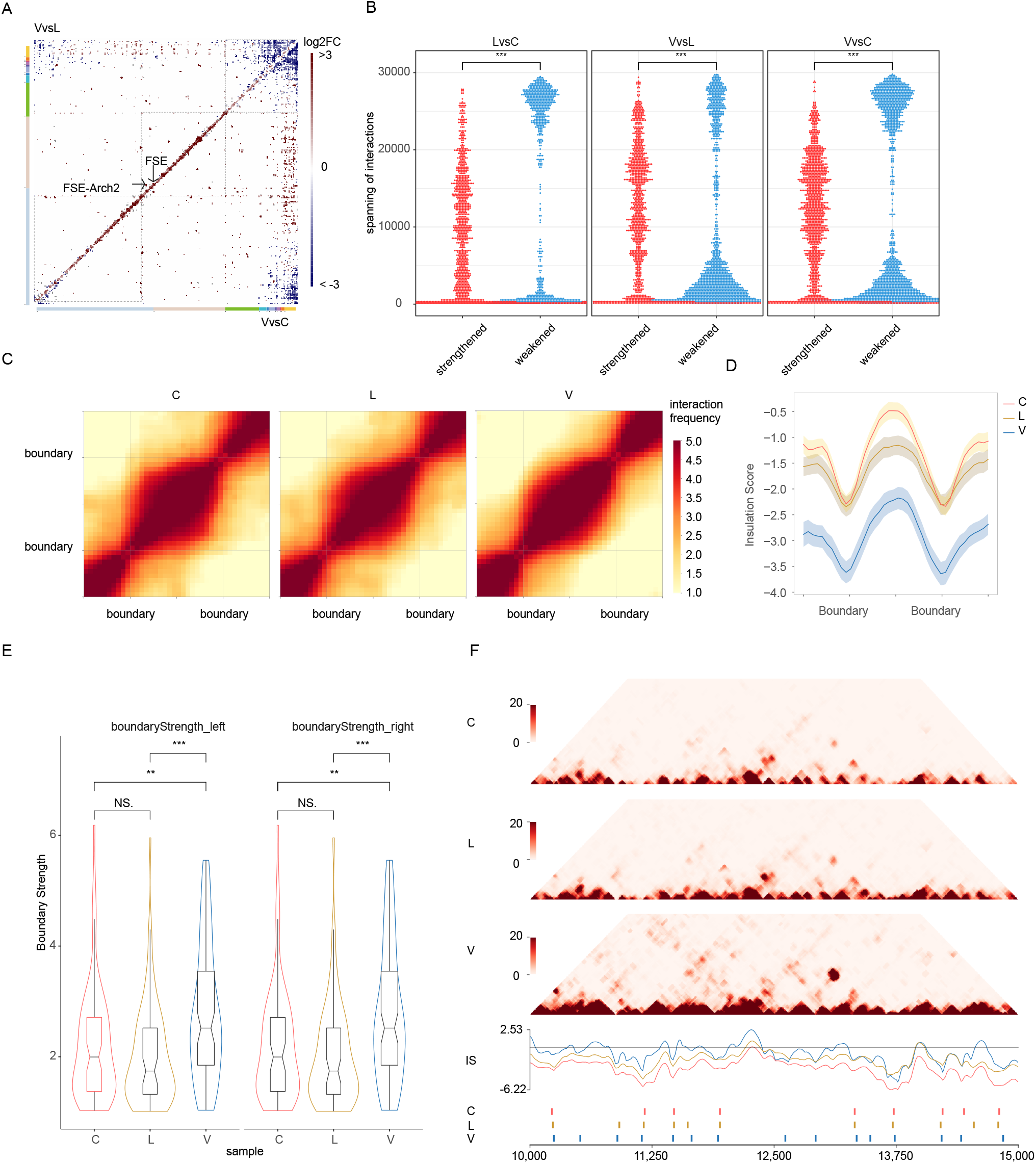
Dynamic structures in different phase of viral cycle. (A) Heatmaps showing comparisons of RNA-RNA interactions in virions vs cells (VvsC) and virions vs lysates (VvsL). VvsL is in the upper quadrant, and VvsC is in the lower quadrant. (B) Different distribution of strengthened and weakened interactions. Dotplots showed distribution of differential interactions as indicated, ***p□<□0.001, two-sided two-sample Kolmogorov–Smirnov test. (C) Maintenance of domains during SARS-CoV-2 virus life cycle. Heatmaps showing the normalized average interaction frequencies for all boundaries as well as their nearby regions (± 0.5 domain length) in C, L and V samples. The heatmaps were binned at 10nt resolution. (D) The average normalized insulation scores were plotted around boundaries from 1/2 domain upstream to 1/2 domain downstream. (E) Violin plot compare boundary strength among C, L and V samples. Showing higher boundary strength in V samples. (F) RNA interaction maps (Top) binned at 10 nt resolution show interactions 10-15kb apart on SARS-CoV-2 genome in C, L and V samples. Line plots (median) show insulation profiles. Short lines (Bottom) reflect boundaries.

We then focused on the changes in interaction mediated by TRS-L region (first 100nt). These interactions were weakened in virions, compared to cells or lysates, both for TRS-L-sgRNA long range interaction and local folding at the `5’`-end of ORF1ab. However, the TRS-L-S interaction was stronger in the lysate than in cells (Figure S7D), perhaps reflecting the first steps of packaging of the viral genome.

The distributions of differential loop lengths are different between cells, lysates and virions (Kolmogorov-Smirnov test, p<2e-16) (Figure 4B). This prompted us to check if these differential interactions are mediated by different kinds of RNAs. RNA-seq coverage in the `3’` third of the genome, where canonical sgRNAs are located, is significantly higher than in the `5’`-two thirds [3]. Therefore, for simplicity, we considered interactions involving fragments `3’` from nt 21,562 to be mediated by sgRNA, and interactions `5’` from nt 21,562 to be mediated by gRNA. We found that most interactions enriched in virions compared to cells or lysates are mediated by gRNA, while interactions depleted in virions are typically mediated by sgRNAs (Figure S7E). In particular, for the differential interactions with long (1kb - 20kb) loops, the weakened interactions are almost always mediated by sgRNAs (Figure S7F). This phenomenon can also be observed in L group compared to C group, indicating that changes in RNA interaction and conformation occur gradually during the virus packaging process. The depletion of interactions mediated by sgRNAs might reflect a decrease in abundance of these sgRNAs. Disruption of long range interaction was also observed for Zika virus inside cells [33], here in SARS-CoV-2, we demonstrated that SARS-CoV-2 RNAs undergoes genome compacting and decreasing in sgRNAs when packaging into virions.

The simplified SPLASH data heatmaps are similar to mammalian genome Hi-C data [34] (Figure 4F), and previous studies suggested that the Zika virus genome is compartmentalized into boundary-demarcated domains [35]. This prompted us to check if SARS-CoV-2 genome RNA are also compartmentalized into domains, and whether global compaction of genome RNA results in impairment of domains. To this end, we applied an insulation score algorithm to call domain boundaries in SARS-CoV-2 genome [36]. In this way, SARS-CoV-2 genome was split into 76, 71 and 96 domains in C, L and V samples, respectively. The average intra-domain contact matrices were shown as heatmap in Figure 4C, indicating a reduction of inter-domain interactions. As expected the insulation scores are significantly lower in boundaries (Figure 4D). Remarkably, the insulation scores are highly correlated between groups of samples (Fig S8A), and domain boundaries are consistent in different samples (Figure 4F). Concordantly, the domain length was comparable between samples (Fig S8 B).

Furthermore, the boundary strength in V are significantly higher than in C and L samples (Figure 4E), and ratio of intra-domain to inter-domain interactions was also higher in V samples (Figure S7C), indicating that during compaction and packaging of the genome, the domain structures were not only retained but even strengthened. Previous studies have revealed domains in Zika virus genome RNA [35] and compaction in virions [33]. Here we described that domains in SARS-COV-2 are stably maintained during life cycle.

Finally, we calculated Shannon entropy values along the SARS-CoV-2 genome (Figure S9A). High entropy indicates flexible regions that may form multiple alternative base-pairs [28]. As expected, the entropies are higher in virions than in cells, indicating that RNAs in cells adopt more alternative structures than in virions (Figure S9B). The entropies inversely correlate with insulation score (Figure S9C), indicating that domain boundaries are more flexible, and might be more attractive sites for drug design.

## Discussion

In this study, we developed a simplified SPLASH protocol based on proximity ligation to capture RNA-RNA interactions in the SARS-CoV-2 virus. RNA proximity ligation has previously been used to address many questions in models ranging from viruses to animal tissues [17, 19, 29, 30, 35, 37-39]. Whereas most previous methods included steps to enrich cross-linked or ligated RNA duplexes, considering low input of RNA amount in virions, we decided to omit all the enrichment steps and hence reduce the complexity of the whole protocol to increase the RNA yield adequate for lib construction and with high depth sequencing. Also, we used RNaseIII to fragment RNA, this made all the treated RNA suitable for RNA ligation, and is supposed to increase efficiency of ligation. As a result, we obtained 28.9% chimeric reads in virions, and more than 10% chimeras in cells and lysates. The higher chimeric rates in virions might result from highly compact genome.

Our results provide the first direct evidence that TRS-L regions form long-distance interactions with TRS-B regions. Although it was widely believed that long range interactions are required for discontinuous transcription, direct experimental evidence for such interactions was lacking. By comparing TRS-L: TRS-B chimeras ligated in `3’`-`5’` and `5’`-3`’` orientations, we distinguished two classes of reads, which represented (1) RNA-RNA interactions, and (2) sgRNAs. We validated two of the putative sgRNAs by Sanger sequencing. Interestingly, although these novel sgRNAs don’t have canonical cCS-B motif upstream of gene body, they both partially overlap with the canonical CS motif. This indicates that TRS-L and at least partial cCS-B base pairing at negative strand are critical for template switching in discontinuous transcription of these sgRNAs. It would be interesting to further identify novel TRS-L dependent sgRNAs and check if they play roles in SARS-CoV-2 biology.

The FSE structure has attracted attention because it is of vital importance in translating nonstructural proteins in ORF1B, and perturbing FSE have significance in modulating coronavirus [40]. A three-stemmed mRNA pseudoknot in the SARS coronavirus frameshift signal was proposed [10] to regulate this process, and was confirmed in SARS-CoV-2 [11]. However, in other cases, distinct structures other than the three-stem pseudoknot were reported [14, 15]. Ziv et al found that the FSE of SARS-CoV-2 is embedded within a ∼1.5 kb long higher-order structure that bridges the 3′ end of ORF1a with the 5′ region of ORF1b, which was termed the FSE-arch [27]. Here we speculate that Ziv’s arch coexists with alternative structures, suggesting that regions around FSE are dynamic, and that RNA conformation changes, presumably to fine tune frameshifting rates and stoichiometry of nonstructural proteins. These additional structures cooperate with Ziv’s arch to embed the FSE in a larger “high order pseudoknot”. The large and small form of pseudoknot might provide a structural basis for ribosome stalling. The mechanisms balancing alternative structures and transcription and translation of orf1a/b remain to be elucidated in the future. Interestingly, the alternative structures described above are also found in virions (Figure 3A), suggesting that even packed into particles, the complicated conformations remained.

By comparing dynamics of SARS-CoV-2 during its life cycle, we found comprehensive compaction of the SARS-CoV-2 genome in virions compared to cells (Figure 4), while genome cyclization and sgRNA-mediated interactions are reduced in virions, and specific TRS-L: S interactions are stronger in virions and in late infected cells (Figure S7D). We found that the genome of SARS-CoV-2 is demarcated by domains, with intra-domain interactions stronger than inter-domain interactions. The uniform and regular domain folding is reminiscent of the nucleosomes-like beaded structure of eukaryotic genome. Importantly, we also found that domains are stronger in virions, and positions of domain boundaries remained consistent during life cycle, with a few domains merged in cells (Figure 4F). Domains were previously reported in the Zika virus [35]. We therefore speculate that domain organization is the rule rather than an exception of genome folding in single strand RNA viruses. Since boundaries remain stable in different phases of virus life cycle, we hypothesize that nucleocapsid (N) protein, packaging RNAs in particle, maintains its role as the regulator of the genome structure in infected cells, when the RNA is released.

---

**Figure S1.**
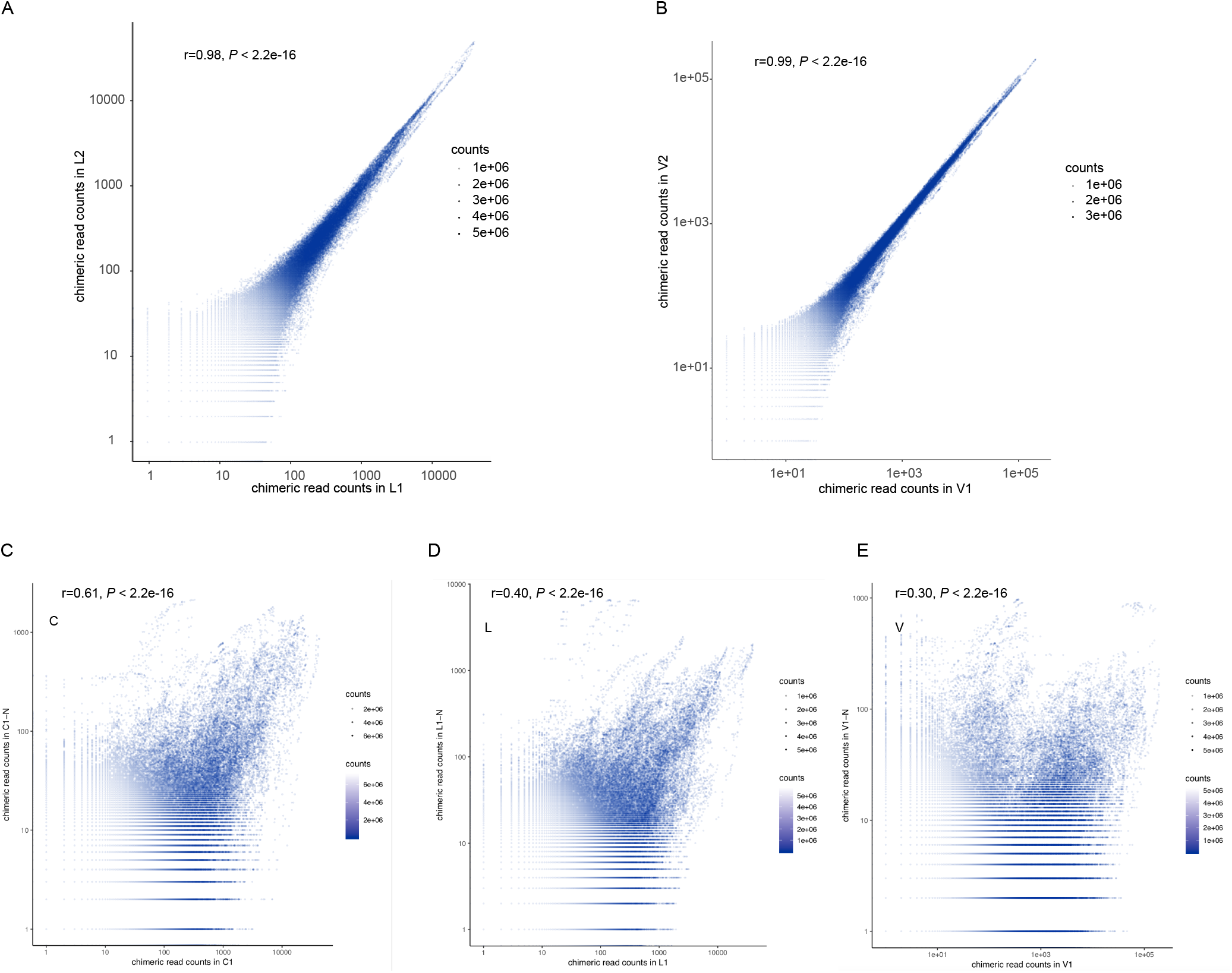
Correlation of chimeric read counts between samples. Scatter plots show correlation of chimeric read counts between ligated samples (A and B), as well as ligated and non-ligated samples (B, C and D).

**Figure S2.**
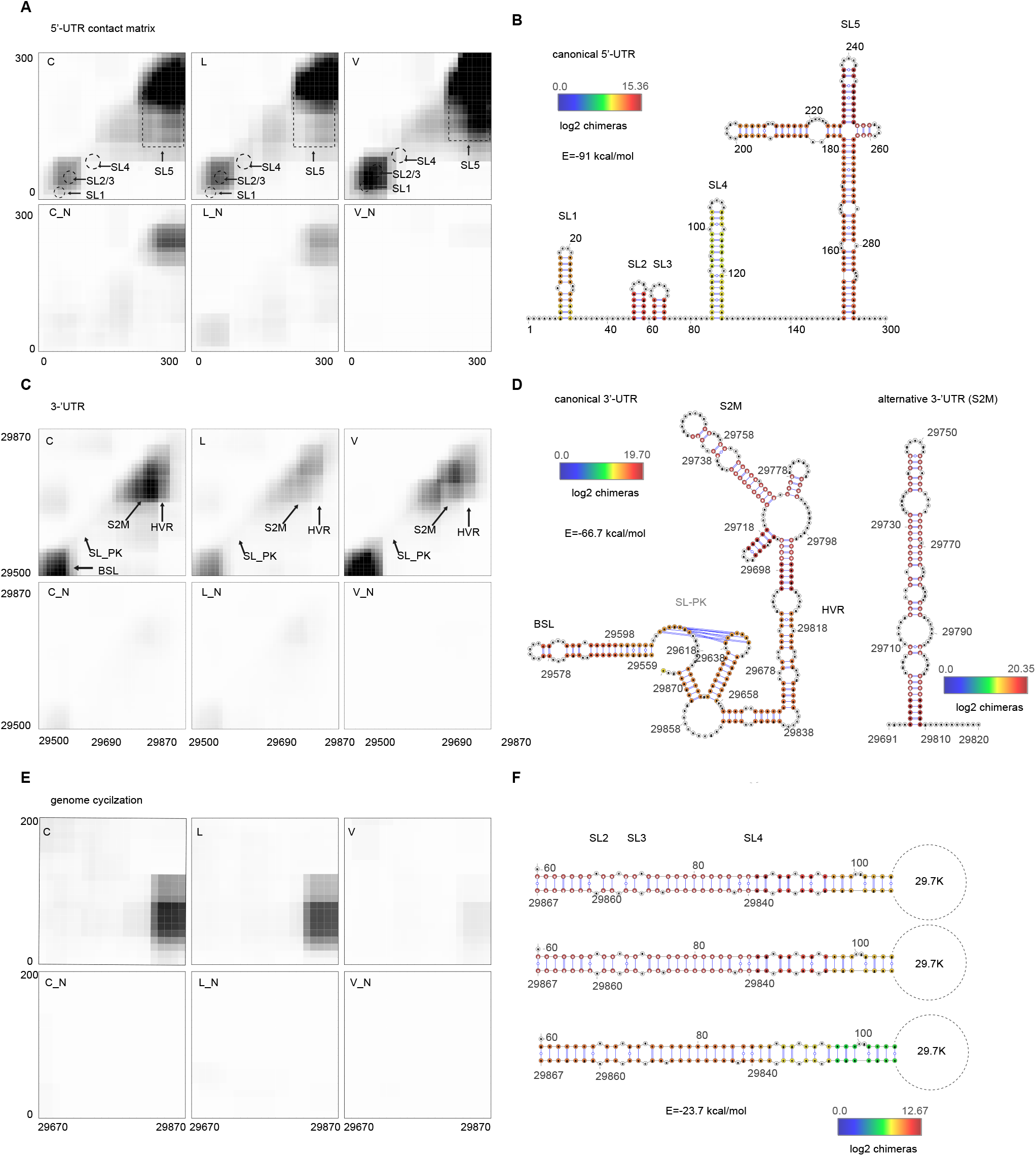
Alternative structures of UTRs. (A) CPM contact matrix in `5’`-UTR region. (B) Canonical SARS-CoV-2 `5’`-UTR structure. Basepairing of indicated arches. Colors represent the log2 chimeric read counts of non-redundant chimeric reads supporting each base-pair. (A) CPM contact matrix in `3’`-UTR region. (D) Canonical SARS-CoV-2 `3’`-UTR and alternative S2M structure. Basepairing of indicated arches. Colors represent the log2 chimeric read counts of non-redundant chimeric reads supporting each base-pair. (E) CPM contact matrix supporting genome cyclization. (F) Base pairing `5’`-UTR and `3’`-UTR in C, L and V samples. Colors represent the log2 chimeric read counts of non-redundant chimeric reads supporting each base-pair.

**Figure S3.**
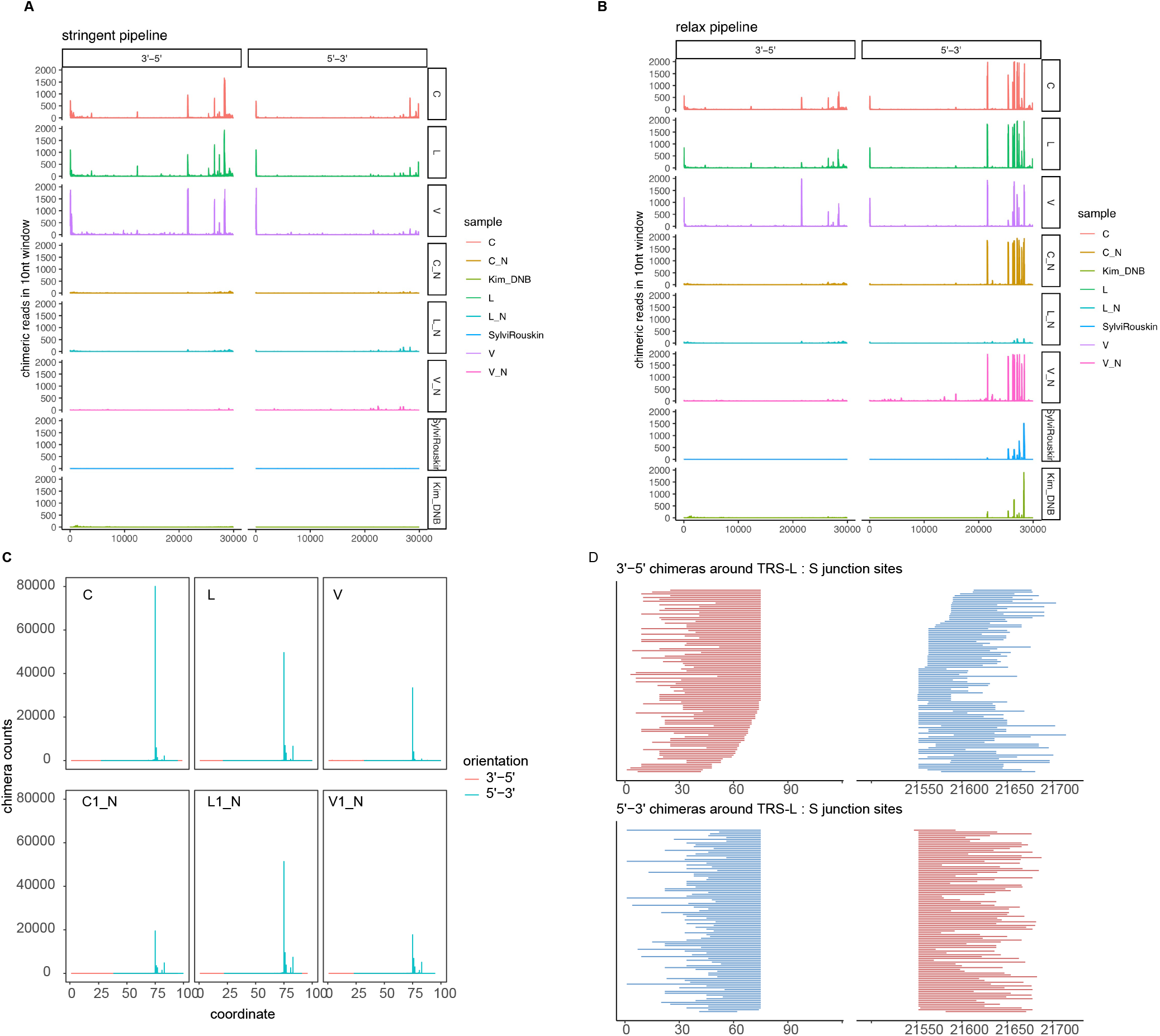
characteristics of TRS-L interaction. (A and B) Viewpoint histograms showing binding positions of the TRS-L region (first 100nt) along the SARS-CoV-2 genome in indicated samples for stringent (A) and relax pipeline(B). `3’`-`5’`chimeras and `5’`-`3’` chimeras were separately plotted. As controls, we also plotted interaction position in non-ligated samples and in RNA-seq data. To show that the interaction peaks especially from `3’`-5 chimeras are specific. (C) Same as Figure 2C, but the chimeras are identified by relax pipeline. (D) Same as Figure 2E, but the chimeras are identified by relax pipeline, which results in plenty of `5’`-`3’` chimeras, so `5’`-`3’` chimeras were also showed, indiccacting highly consensus of junction sites.

**Figure S4.**
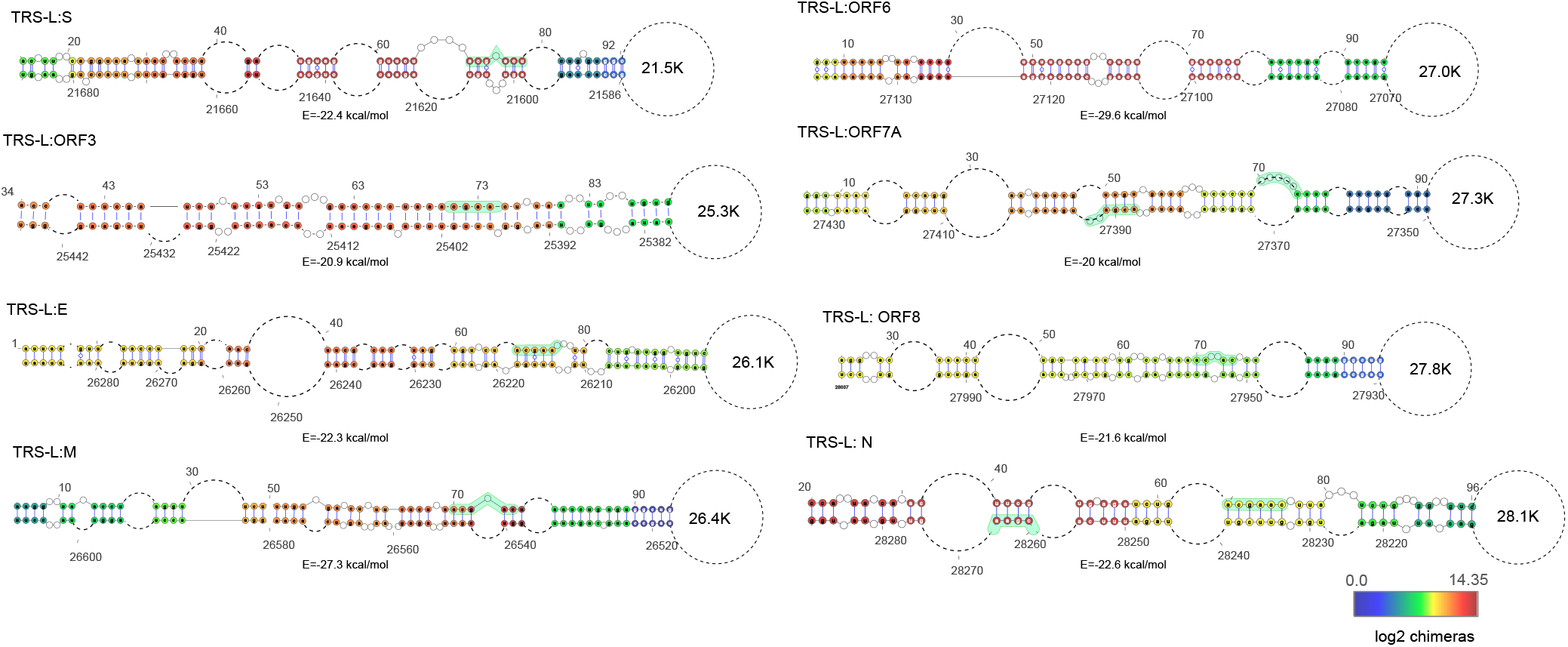
Characteristics of TRS-L interaction. RNA secondary structures TRS-L and TRS-B interactions identified from this study.

**Figure S5.**
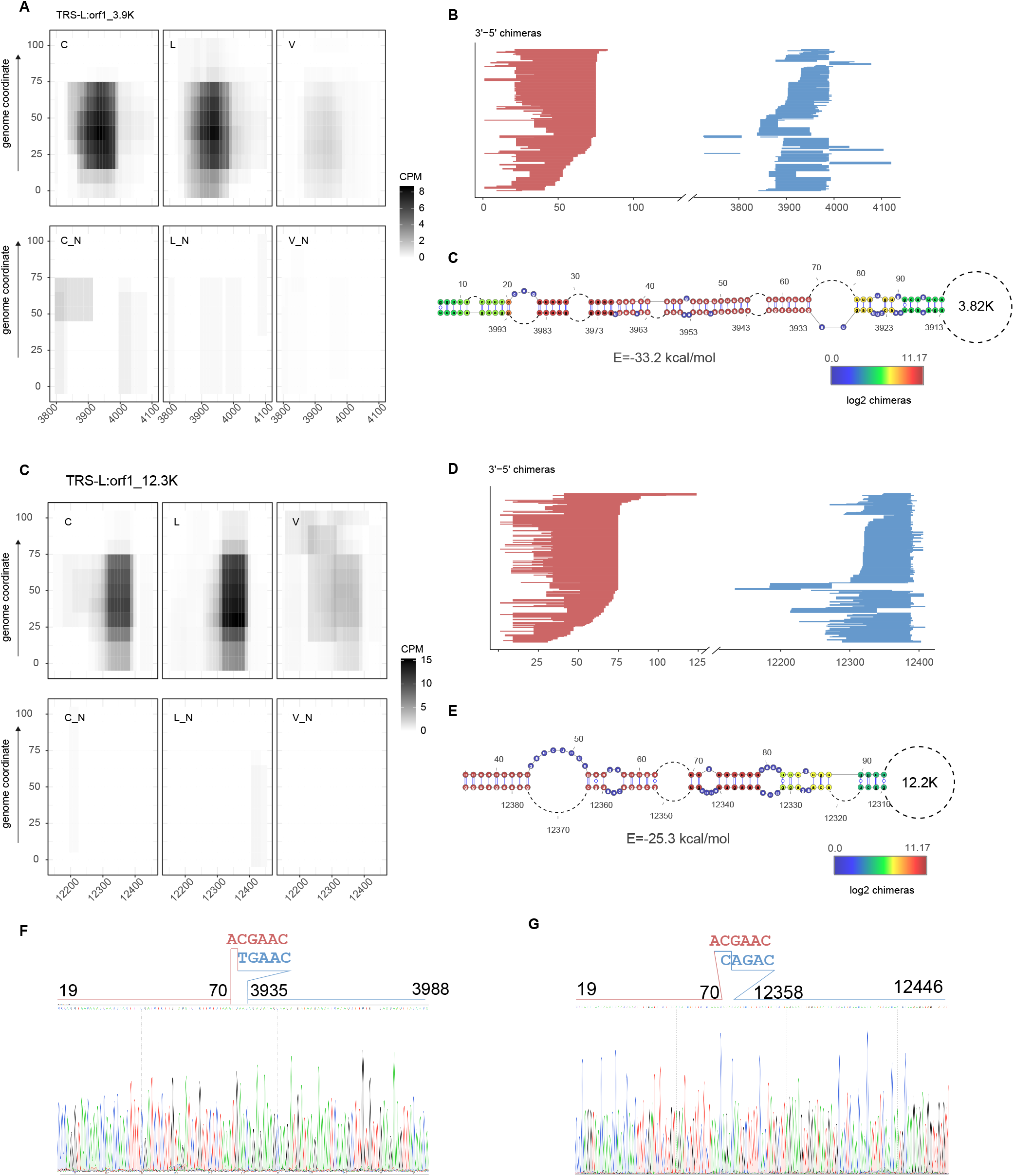
Identification and validation of novel sgRNAs. (A and C) Contact matrix of `3’`-5 chimeric reads indicated specific interactions of TRS-L and 3.9K (A) and 12.3K (regions). Color depict chimeric reads per 1million mapped reads (CPM). (B and D) Randomly selected chimeric reads showed distribution of reads aounrd junction sites. The redlines indicated `3’`arm in chimeric reads, while blue lines indicated `5’`arrm in chimeric reads. (C and E) base pairing of TRS-L region and 3.9k (C) and 12.3K(E) regions (F and G) Sanger sequencing validation of 3.9K (F) and 12.3K(G) novel sgRNAs from independent non-crosslinked samples. Bases around junction sites were shown.

**Figure S6.**
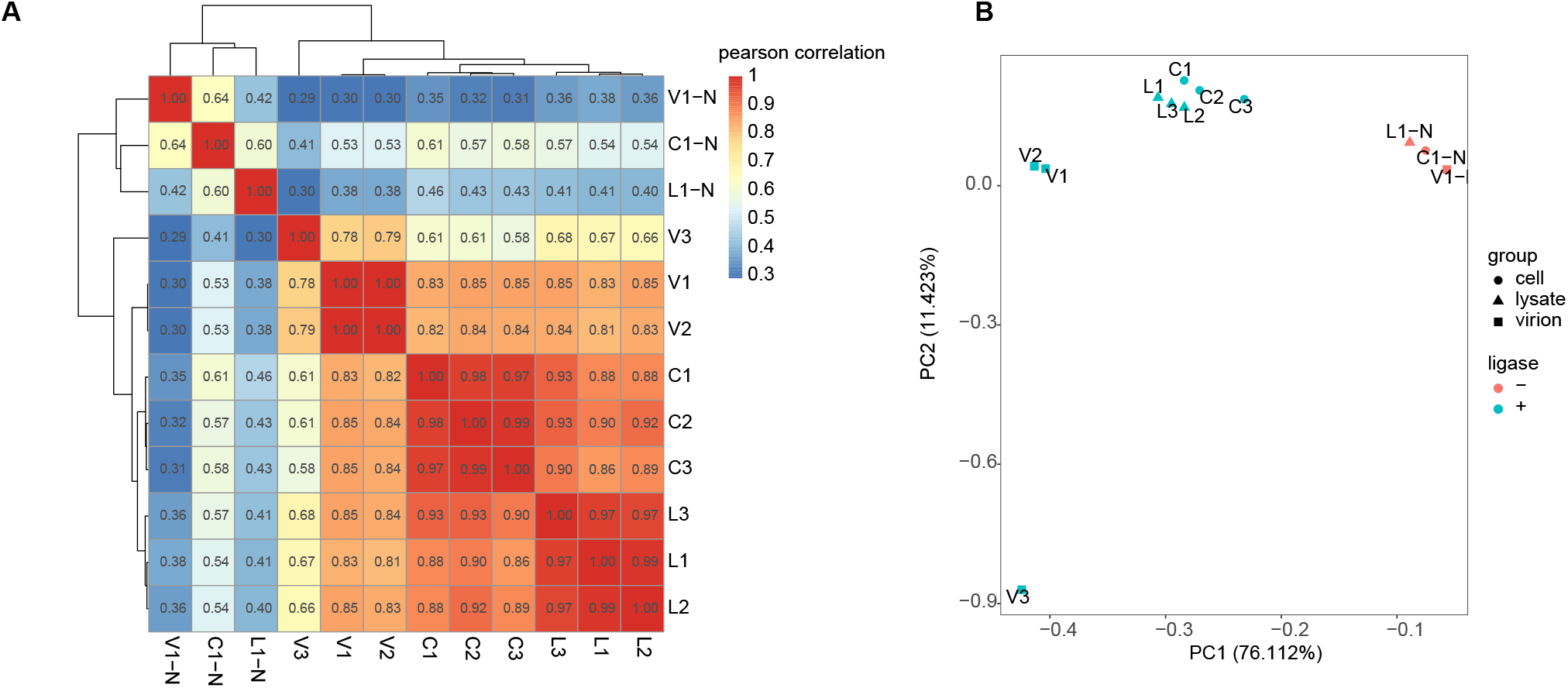
Clustering and PCA of samples. (A) cluster of Pearson correlation efficiencies between samples. (B) Principal component analysis (PCA) interaction data from samples indicated in A.

**Figure S7.**
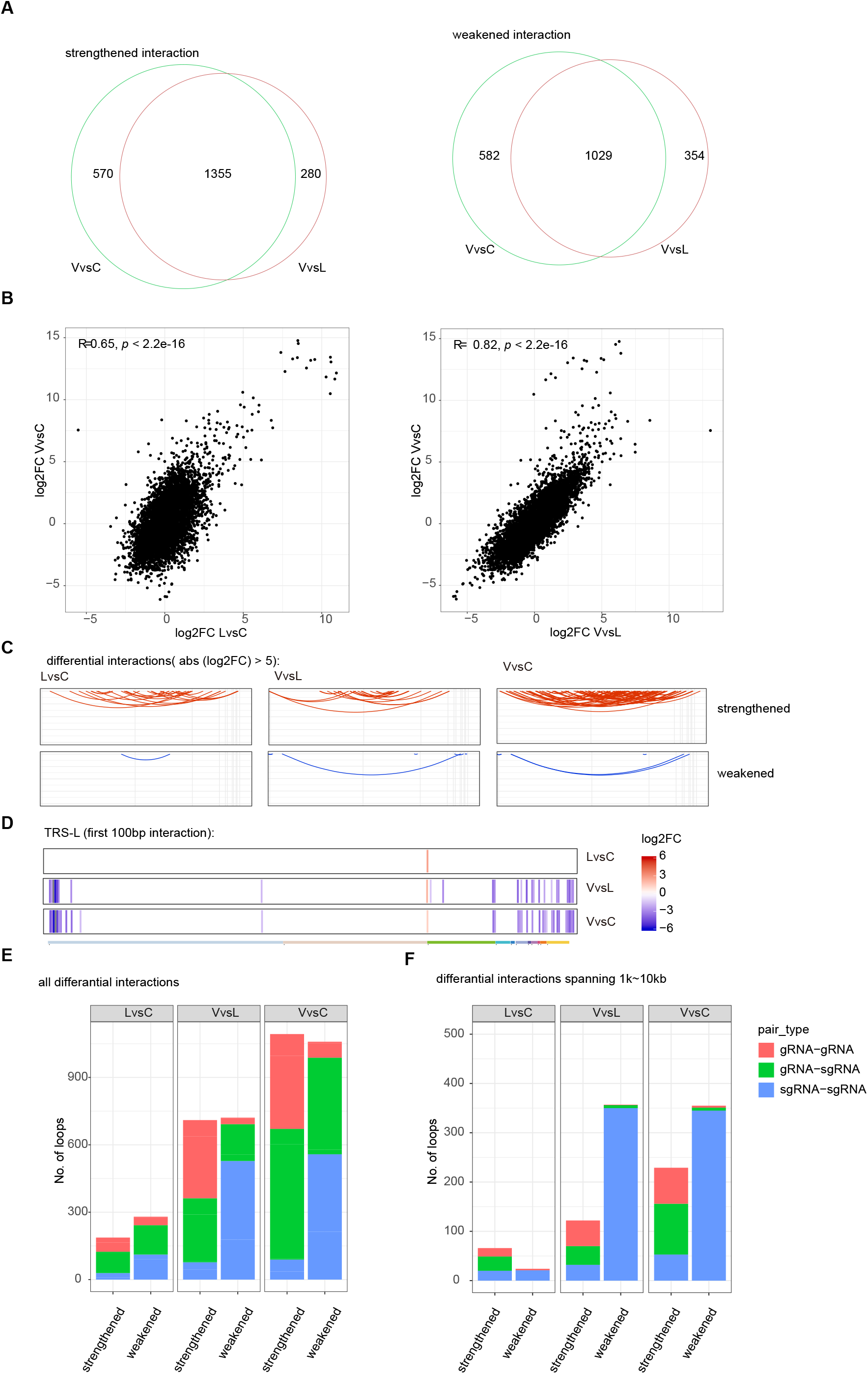
Dynamics of interactions in different phase of SARS-CoV-2 virus life cycle. (A) Vennplot overlap of differential interactions of VvsC and VvsL. (B) correlation of log2FCs as indicated. (C) Arc plots show strengthened and weakened interactions. Differential interactions with abs(log2FC) cutoff >= ±5 were plotted (D) Heatmaps show log2FC of all the TRS-L (first 100nt) interactions in different comparisons. Note that almost all interactions are not changed or weakened, except that TRS-L: S interaction is strengthened. (E) RNA type distribution of differential interactions. For simplicity, RNA position after nt21562 are considered as sgRNAs, and considered as gRNA before nt21562. (F) Same as (E), except that only differential interactions spanning from 1kb to 10kb were analyzed.

**Figure S8.**
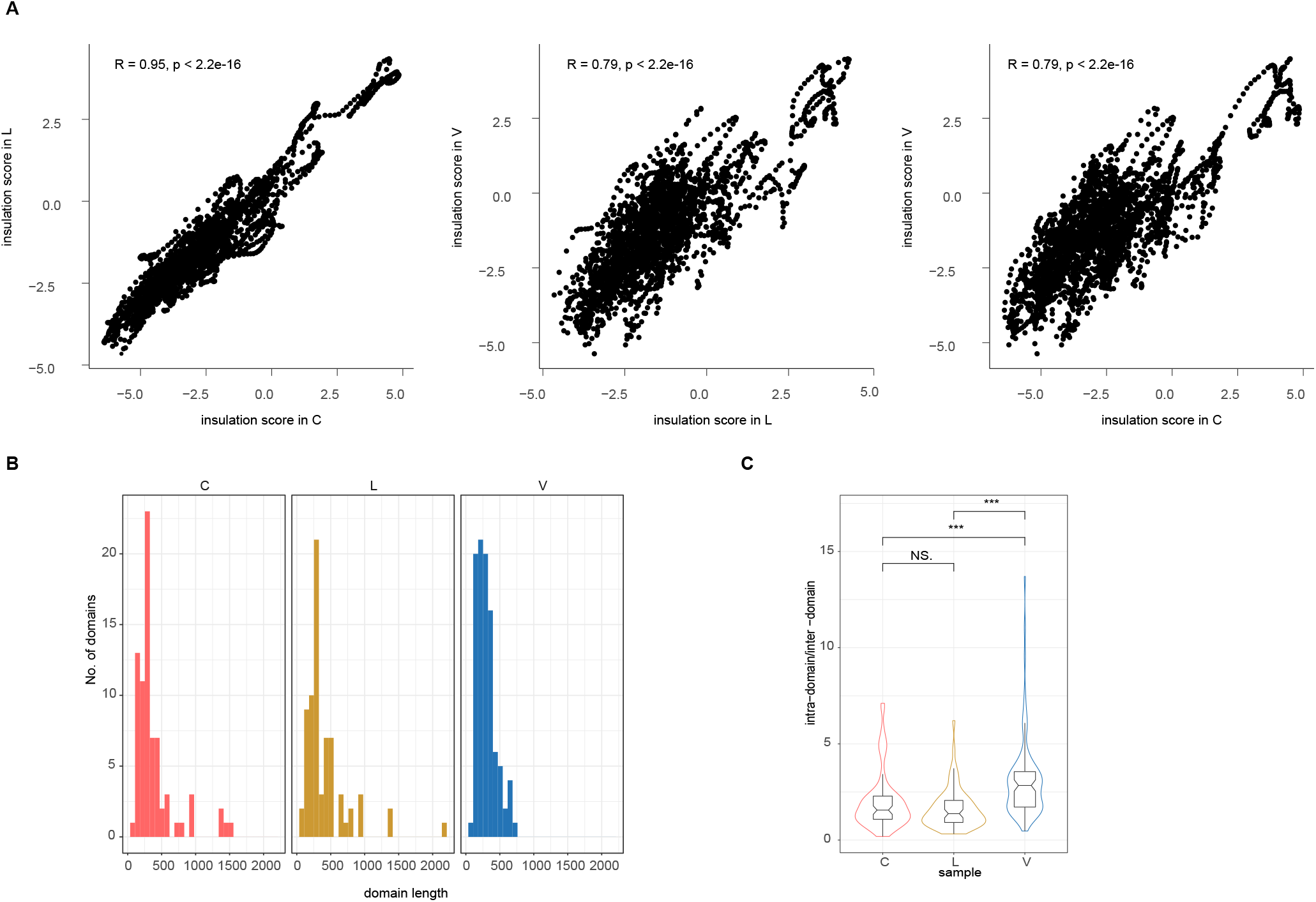
Characteristics of boundaries in different phase of SARS-CoV-2 virus life cycle. (A) Correlation of insulation score in different samples. (B) Histograms of boundary length in different samples. (C) Violin plot compare ratio of intra-domain/inter-domain interactions.

**Figure S9.**
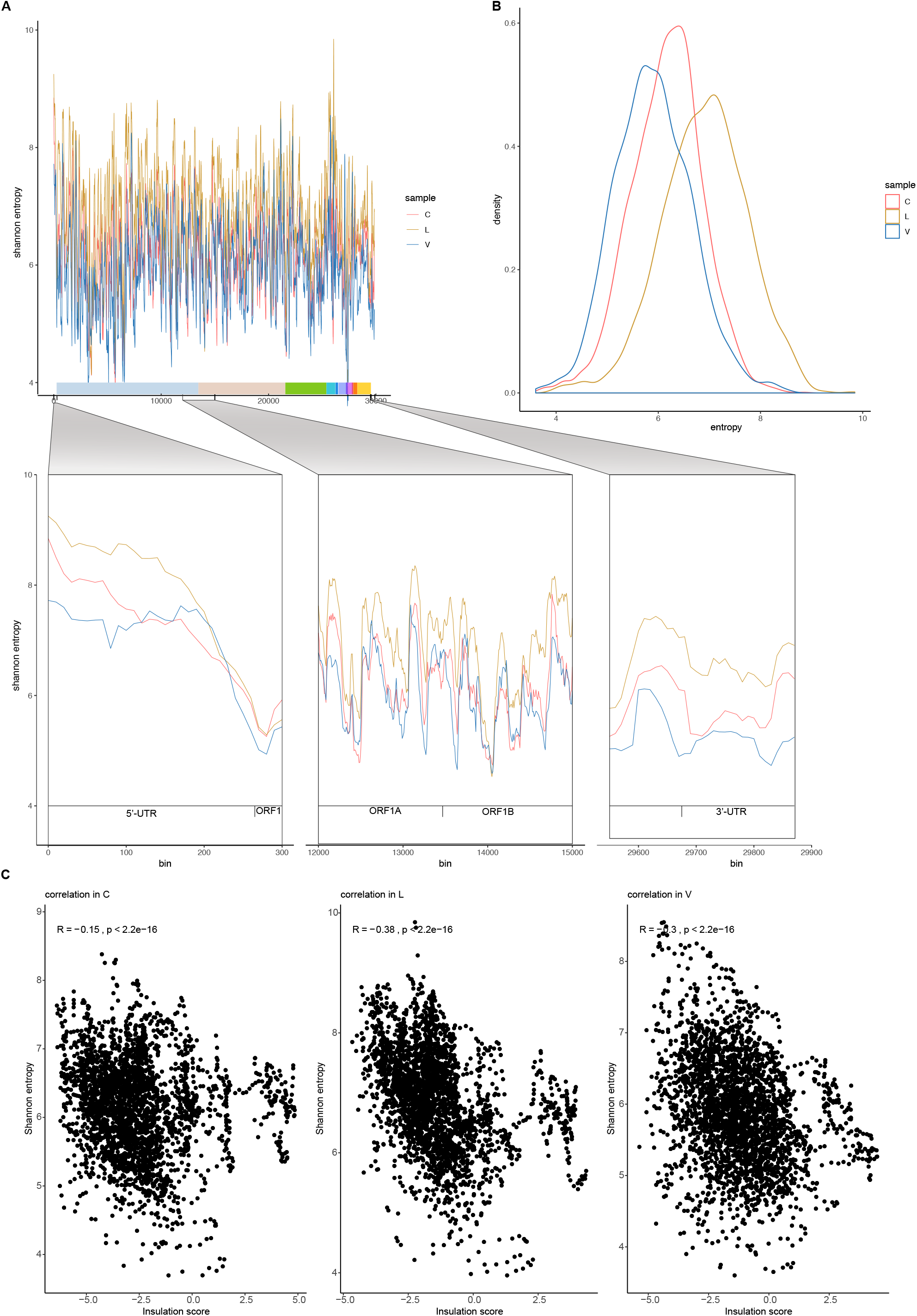
Shannon entropy of SARS-CoV-2 genome. (A) Shannon entropy values along the SARS-CoV-2 genome. (B) density plot of Shannon entropies from different samples. Shannon entropies were highest in L while lowest in V, tested by wilcox test (P<2.2e-16) (C), Shannon entropy values for selected regions

## Supplementary table1

Statistics of mapping and chimeras of all the samples

## Materials and Methods

### Cell lines

Chlorocebus sabaeus (Green monkey) VeroE6 (female, RRID:CVCL_YQ49) were purchased from American Type Culture Collection (ATCC, id: ATCC CRL-1586). Vero E6 and VeroE6/TMPRSS2 cells were cultured in Dulbecco’s modified Eagles medium (DMEM) supplemented with 10% fetal bovine serum at 37 °C in a humidified CO2 incubator.

### Virus inoculation and crosslinking

Infection experiments were performed under biosafety level 3 conditions. SARS-CoV-2 virus strain Wuhan-Hu-1 was kindly provided by (Wuhan institute of viology). Independent biological replicates were performed using 90-120 million cells each. VeroE6 cells were inoculated with SARS-CoV-2 strain Wuhan-Hu-1 at MOI=0.01 pfu/cell for 24 hours. Following inoculation, two flasks of cells were washed 3 times by PBS and then subjected to crosslinking. The remaining cells were cultured for another 48hours, when cytopathic effect (CPE) was observed in about 70% cells, supernatant was collected and centrifuged at 4°C 1000rpm for 10min to remove cell pellet. Then the clear supernatant was mixed with equal volume of saturated ammonium sulfate and incubate at 4°C for one hour. At the same time, remaining unshed cells were washed 3 times by PBS and subjected to crosslinking.

For crosslinking, cells or virus pellet were incubated with 2 mM of EZ-Link Psoralen-PEG3-Biotin (ThermoFisher Scientific) at 37°C for 10 min in PBS containing 0.01% digitonin. The cells were then spread onto a 10cm plate and irradiated using 365 nm UV for 20 min on ice. Cell and virion RNA were extracted with RNeasy mini kit (Qiagen).

### simplified SPLASH assay

500 ng of RNA was fragmented using RNase III (Ambion) in 20μl mixture for 10miniutes at 37°C, and purified using 40 μl of MagicPure RNA Beads (TransGen). Each RNA sample was divided in two: one half was used for proximity ligation and then crosslink reversal (C, L and V samples), while in the other half, crosslink reversal was done before proximity ligation (non-ligated C_N, L_N, V_N). Proximity ligation was done under the following conditions: 200ng fragmented RNA, 1 unit/μl RNA ligase 1 (New England Biolabs), 1× RNA ligase buffer, 50 mM ATP, 1 unit/μl Superase-in (Invitrogen), final volume: 200 μl. Reactions were incubated for 16 hours at 16 °C and were terminated by cleaning with miRNeay kit (Qiagen). Crosslink reversal was done by irradiating the RNA on ice 254 nm UVC for 5min using a CL-1000 crosslinker (UVP).

### Sequencing library preparation

Sequencing libraries were prepared with 50ng input RNA material using SMARTer Stranded Total RNA-seq Kit v2—Pico Input Mammalian (Takara Bio USA Inc., USA), according to the manufacturer’s instructions. The libraries were paired-end sequenced (PE150) using Illumina Nova seq platform.

### Data pre-processing

Data preprocessing was performed according to (Ziv et al., 2018). In brief, raw paired-end reads were trimmed for adaptors and checked for quality using cutadapt (Martin, 2011). Chimeric reads were identified and annotated to the respective genome using hyb (Travis et al., 2014). SARS-CoV2 samples were processed using SARS-CoV-2 sequence (NC_045512.2).

### Chimeras and interaction calling

Chimeric reads were called and annotated with the hyb package [41], using the command:

default (stringent) parameters:

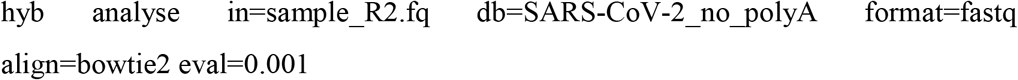

Relaxed pipeline:

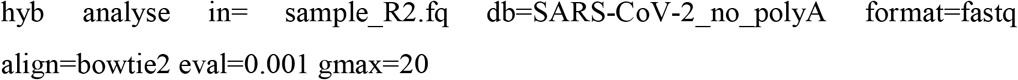

To evaluate the folding energy of chimeric reads, we used hybrid-min28 with default settings. We then randomly reassigned (shuffled) pairs of fragments found in chimeric reads, and repeated the folding energy analysis. The folding energies of experimentally identified and shuffled chimeras were compared by Wilcoxon test.

Virus interaction heatmaps were plotted using Java Treeview [42], as previously described [28], such that color intensity represents the coverage of chimeric reads at every pair of positions. The first read of each pair is plotted along the X axis, and the second read along the Y axis. As a result, chimeras found in the 5’-3’ orientation are shown above the diagonal, and chimeras in the 3’-5’ orientation are below the diagonal. Viewpoint histograms and arc plots were plotted with ggplot2 R package [43].

For TRS-L interaction peak calling, each chimeric read was split and mapped to two paired non-overlapping 10nt-bins, we first scored interactions by log2 transformed chimeric reads, then calculated z-scores for all the interaction pairs. z-scores > 2.13 (which means log2 chimeras larger than average with 95% confidential) were considered as enriched interactions. Then enriched TRS-L interaction were selected if either arm located in 1-100nt.

### RNA secondary structure folding

For short range interactions, we assembled non-zero chimeric groups into uninterrupted stem structures, and then fold RNA using COMRADES (https://github.com/gkudla/comrades).

For long range interactions, we first assembled uninterrupted stem structures as above for each arm, and then fold RNA by hybrid-min in unafold-3.8 [44].

Folded structures were visualized in VARNAv3-93.

### Calling of Topological Domain

Domain boundaries were identified by insulation score [36] using the 10 nt resolution simplified SPLASH contact matrices data. Here we used 500 nt×500 nt (50× resolution) square along each bin for calculating insulation score, A 150 nt (10× resolution) window was used for statistics of the delta vector and removed the weak boundaries which ‘boundary strength’ lower than 1.

### Average Insulation Score of Domain

The average insulation scores were normalized by around all domain as well as their nearby regions (± 0.5 domain length). The heatmaps were binned at 10 nt resolution and an 800 nt window. Average insulation scores were plotted around boundaries from 1/2 domain upstream to 1/2 domain downstream.

### Average interaction heatmap of domains

The size of the domain was homogenized to 400 nt, the upstream and downstream extended 1 / 2 domain. calculated the interaction frequency by the averaging all domain. The resulted matrices were plotted as heatmap by log2 average signals.

### 3D modeling of virus genome

We used pastis-0.1.0[45] software to model RNA genome in three dimensions. The final results obtained by MDS algorithm were used for 3D visualization. For the spatial location of particular gene loci, we used 1 point/20 balls to calculate the position of specific genes in the whole 3D simulation, and then modify the pymol results by a python script.

### Differential interaction identifying

Differential gene expression analysis was performed using DESeq2 [32]. 100nt-Bin interactions that displayed more than ±1 log2FC (FDR <0.01) between C or L and V samples were considered as significantly differential interactions. Then log2FC heatmaps were plotted in using Java Treeview.

### RT-PCR of candidate novel sgRNAs

RNA from infected non-crosslinked VERO celles (24hours, as described above) were extracted by miRNeasy Mini Kit (Qiagen). 100ng of RNA was then subjected to retrotranscription using SuperScript VILO Master Mix (Thermo Fisher), cDNA was amplified with 2xEs Taq MasterMix (Cowin Biotech) by 0.4 μM of each primer: 3.9K (TGTTGTAACTTCTTCAACACAAGC) or 12.3K (TGTTCAAGGGAACACAACCATC) and TRS-L (CCCAGGTAACAAACCAACCAAC).

## Data availability

Raw and processed sequencing datasets analyzed in this study have been deposited in the Gene Expression Omnibus (GEO) database, https://www.ncbi.nlm.nih.gov/geo/ (accession number GEO: GSE164565).

## Statistics

Statistical analyses for differential interaction was conducted with the R Bioconductor package DESeq2 using three independent replicates as described above.

Comparing quantitative indicators such as boundaries strength, was performed with two-sided Wilcox rank sum test.

To test whether differential interactions spannings follow the same continuous distribution, two-sided Kolmogorov-Smirnov test was performed.

Statistical significance of differences in odds ratios between two groups (Figure S6E, S6F) was calculated using a two-sided Fisher`’`s exact test.

Correlation analysis of chimeric reads counts (Figure 1B, S1) and insulation scores (Figure S7A) between samples were performed by Pearson’s product moment correlation coefficient (R or PCC).

All the statistics tests were performed with R package stats.

## Author Contributions

Z. Zhao conceived and supervised the project, G. K supervised design and bioformatic analysi of the project, M. J supervised virus infection and cell culture; Y. Z performed simplified SPLASH experiments, K.H and Z. Zou performed virus and cell culture and crosslinking, D. X, J. L., H. R and W. S. performed bioinformatics analysis, D. W., S. S. and P. L. performed library construction and RT-PCR experiments. Y. Z, Z. Zhao and G.K wrote the manuscript. All authors critically revised the manuscript.

## Conflict of Interest

The authors declare no competing interests.

## Acknowledgment

This study was supported by National science and technology major projects (2018ZX10305410) to Z. Zhao, and National Key Research and Development Program of China (2018YFA0900801) to Y. Z and Wellcome grant 207507 to G. K.

## Notes

### Competing Interest Statement

The authors have declared no competing interest.

### Summary of Updates

The co-first and co-corresponding authors are marked in the revised manuscripts.

